# Ergodic patterns of cell state transitions underlie the reproducibility of embryonic development

**DOI:** 10.1101/2021.08.23.457360

**Authors:** Miriam Genuth, Yasuhiro Kojima, Dörthe Jülich, Hisanori Kiryu, Scott A. Holley

## Abstract

The reproducibility of embryonic development is a remarkable feat of biological organization, but the underlying mechanisms are poorly understood. Clearly, gene regulatory networks are central to the orderly progression of development, but noisy molecular and cellular processes should reduce reproducibility. Here, we identify ergodicity, a type of dynamical stability, as underlying the reproducibility of development. In ergodic systems, a single timepoint measurement equals a time average. Focusing on the zebrafish tailbud, we define gene expression and cell motion states using a parallel statistical analyses of single cell RNA sequencing data and in vivo timelapse cell tracking data and a change point detection algorithm. Strikingly, the cell motion state transitions in each embryo exhibit the same patterns for both a single timepoint and a 2-3 hour time average. Both the cell motion and gene expression cell states exhibit balanced influx and outflux rates reflecting a spatiotemporal stability. Stated simply, these data indicate the pattern of changes in the tailbud doesn’t change. This ergodic pattern of cell state transitions may represent an emergent meta-state that links gene networks to the reproducible progression of embryogenesis.

## Introduction

Aristotle first noted the astonishing reproducibility of embryogenesis in “Historia Animalium,” where he observed, “Generation from the egg occurs in an identical manner in all birds”. At the cellular level, development entails a reproducible series of cell state transitions representing changes in gene expression state, physical state and cell fate. These processes can be noisy, for example, cell migration can be either ordered or disordered, and such disorder is part of normal orderly development. We now appreciate that gene networks control cell state transitions, but these networks are comprised of stochastic molecular processes. How biological order emerges from stochastic molecular events was the subject of Erwin Schrödinger’s “What is Life?”. Despite the remarkable progress in the field of developmental biology in recent decades, there is still a gap in our understanding of the organizing processes that lie between genes and the reproducible dynamics of developing embryos.

One framework for analyzing the reproducibility of embryonic development is that of ergodicity. An ergodic system is one in which the average behavior of all objects at a single timepoint equals the average behavior of a random sample of objects over a longer time interval. For example, measurement of all gas molecules within a chamber at a single time point yields the same result as the average of a random sample of gas molecules over the entire experimental time interval. The ergodic hypothesis lacked a mathematical foundation until the development of the ergodic theorem in 1932. In the field of ergodic theory, the single timepoint average is typically referred to as the “phase average”. Ergodicity is a mathematical ideal and real systems are not truly ergodic. Embryonic development is by definition not ergodic since the embryo changes as it develops, but it is possible for ergodicity to exist over a short period of time. Ergodicity is implicitly assumed in many biological experiments, yet it is rarely demonstrated. For example, ergodicity of gene expression levels in clonal cell populations is only observed when accounting for cell age.

Here, we address the mechanism of the reproducibility of embryonic development by performing an analysis of ergodicity of the pattern of cell state transitions in the zebrafish tailbud. The vertebrate tailbud is a dynamic structure that supports body elongation (Fig. 1a, left panel). Cells in the tailbud undergo multiple transitions in gene expression and migratory behavior during their differentiation. The dorsal-medial tailbud (DM) contains a pool of *sox2/brachyury* expressing neuromesodermal progenitors (NMPs) that contribute to both the spinal cord (Fig. 1A, yellow) and the presomitic mesoderm (PSM). In the zebrafish, cells in the DM migrate towards the posterior in a processive orderly fashion (Fig. 1A, cyan). At the tip of the tailbud, mesodermally fated cells downregulate *sox2*, upregulate mesodermal genes such as *tbx16*, and undergo EMT to migrate ventrally into the progenitor zone (PZ)(Fig. 1A, magenta). Cell movements in the PZ are more disorderly than the DM. Ultimately cells leave the PZ, reduce their cell motion and assimilate into the left and right PSM (Fig. 1A, green). Cells in the PSM downregulate *tbx16* and turn on *tbx6*. Cell velocity in the anterior PSM declines further as the tissue solidifies. The transition from orderly to disorderly motion from the DM to PZ is necessary for proper body elongation. Excessively disordered motion in the DM (obtained by inhibition of BMP or FGF signaling) impairs the flow of cells through the tailbud leading to a short body axis. Excessively ordered motion in the PZ (induced by moderate Wnt inhibition) produces prolonged anisotropic fluxes, unequal allotment of cells to the left or right PSM, and a bent body axis. Thus, understanding robustness and reproducibility of vertebrate body elongation requires understanding the nature of these tailbud cell state transitions.

**Fig. 1.**
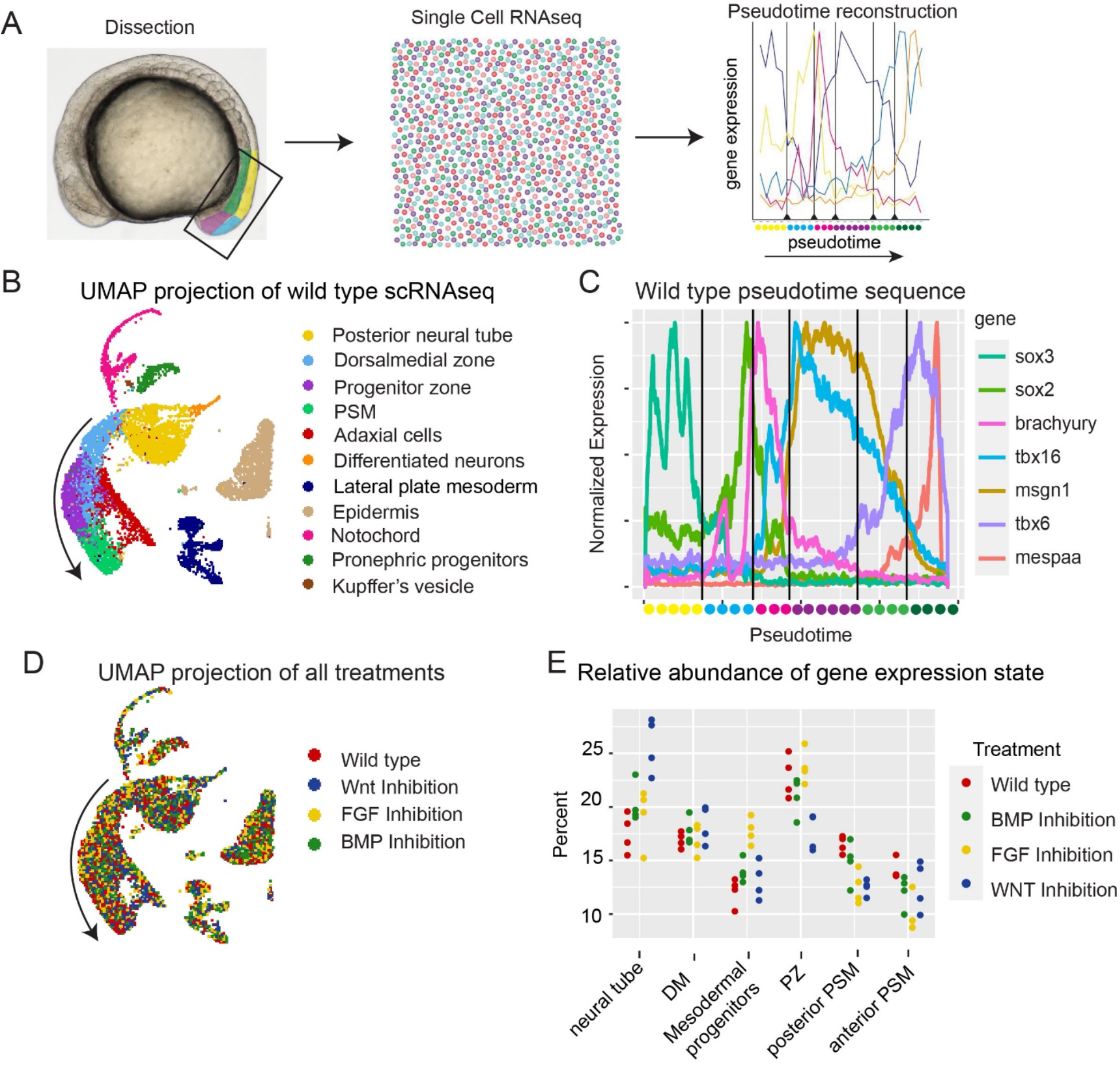
Gene expression cell states in the zebrafish tailbud. **(A)** Schematic of the experimental approach. Tailbuds were dissected and pooled, scRNAseq profiles were generated, and a one-dimensional pseudotime was created and segmented into gene expression cell states. **(B)** UMAP projection of scRNAseq data colored by cell type. Arrow marks the path of pseudotime in C. **(C)** Expression of selected markers over pseudotime. Vertical lines are transition points between cell states as defined by a Bayesian algorithm that minimizes within state statistical error. Note that the segment colors along the pseudotime axis correspond to the colors along the developmental trajectory (arrow) in B but with the Progenitor Zone and PSM being further subdivided into two similarly colored segments in C. **(D)** UMAP projection of scRNAseq data colored by experimental treatment. **(E)** Quantification of the differences in the proportion of cells that are in a given cell state in each replicate of each experimental condition. See also Figs. S1, S2 and Table S1.

In this study of ergodicity and the reproducibility of development, we first objectively define the trajectory of cell states in the zebrafish tailbud during body elongation. We define gene expression state using single cell RNA sequencing (scRNAseq) and a change point detection algorithm. We validate this method by comparing wild type and embryos with reduced Wnt, Fgf and Bmp signaling, and verify quantitative differences in cell states by multicolor fluorescent in situ hybridization. Next, we identify cell motion states by analyzing cell tracking data using the same statistical analysis as used for scRNAseq. We then perform an analysis of ergodicity of the pattern of cell motion states as these datasets allow direct comparison of a single timepoint with a time average. We find that the ergodicity is achieved via a balanced flux between cell motion states in each embryo. Since it is not possible to directly track gene expression states over time in scRNAseq data, we estimate the flux between the gene expression states from RNA velocity and confirmed balanced flux in wild-type embryos. These consistently balanced fluxes for both cell motion and gene expression states suggest that the ergodicity is an emergent order which can explain the reproducibility and robustness of embryonic development.

## Results

### Gene expression states

We performed scRNAseq on dissected tails from 10-12 somite stage zebrafish embryos (Fig. 1A). We used wild-type embryos and embryos subject to treatments known to alter tailbud cell migration, specifically inhibition of FGF, BMP, or Wnt signaling. For each treatment, we prepared four biological replicates each consisting of 10 to 12 tailbuds and resulting in 30,000-35,000 single cell profiles. In a UMAP dimension reduction plot of wild type, the neuronal and paraxial mesoderm form one large cluster with more differentiated cells at each end and common progenitors (cyan) in the middle (Fig. 1B, arrow, and Fig. S1). Wild-type and experimental samples consist of the same cell transcription profiles (Fig. 1D). This result is consistent with previous scRNAseq analysis of zebrafish embryos indicating that perturbation of cell signaling does not create novel cell transcription profiles.

To enable direct quantitative comparisons between experimental conditions, we pooled the data from all wild-type and experimental replicates and created one unified pseudotime to define a single standard for classifying cells. Specifically, the cells in the main cluster were aligned along a neuronal-mesodermal axis from *sox3* expressing neuronal cells to *mespaa* expressing anterior PSM cells (Fig. 1B, arrow). This approach avoids the requirement to define the NMP population a priori. Instead, NMPs will be located in the middle of the pseudotime sequence and differentiation will proceed towards both ends, i.e. neuronal to the left and mesodermal to the right (Fig. 1C). Marker genes for neuronal and mesodermal development map with respect to pseudotime in the correct developmental sequence indicating that the procedure was successful.

To objectively define gene expression states, we extracted the wild-type data, and then utilized a change point detection algorithm to divide pseudotime into a series of distinct states. The change point algorithm identified five transition points (Fig. S2). These transition points (vertical lines in Fig. 1C) divide the pseudotime sequence into six states that generally agree with those predicted previously from marker gene expression. These transition points were mapped to the full pseudotime sequence, and we calculated the relative abundance of each state in wild type, Wnt inhibited embryos, Fgf inhibited embryos and Bmp inhibited embryos (Fig. 1D).

To determine whether this analysis of scRNAseq data accurately quantifies changes in cell state, we mapped the transcriptional states back onto the embryo and measured their abundance using simultaneous multicolor fluorescent in situ hybridization for marker genes for the first five states (Fig. 2A). *Sox2* single positive cells localize in the neural tube (state 1). *Sox2* and *brachyury* positive NMPs (state 2) occupy the DM. Nascent mesodermal progenitors (state 3) expressing *brachyury* and *tbx16* are located immediately ventral to the DM in the medial PZ. Mesodermal progenitors in the PZ (state 4) are *tbx16* single positive cells located in the ventral and lateral tailbud. The PSM (state 5) is anterior to the transition from *tbx16* to *tbx6* expression.

**Fig. 2.**
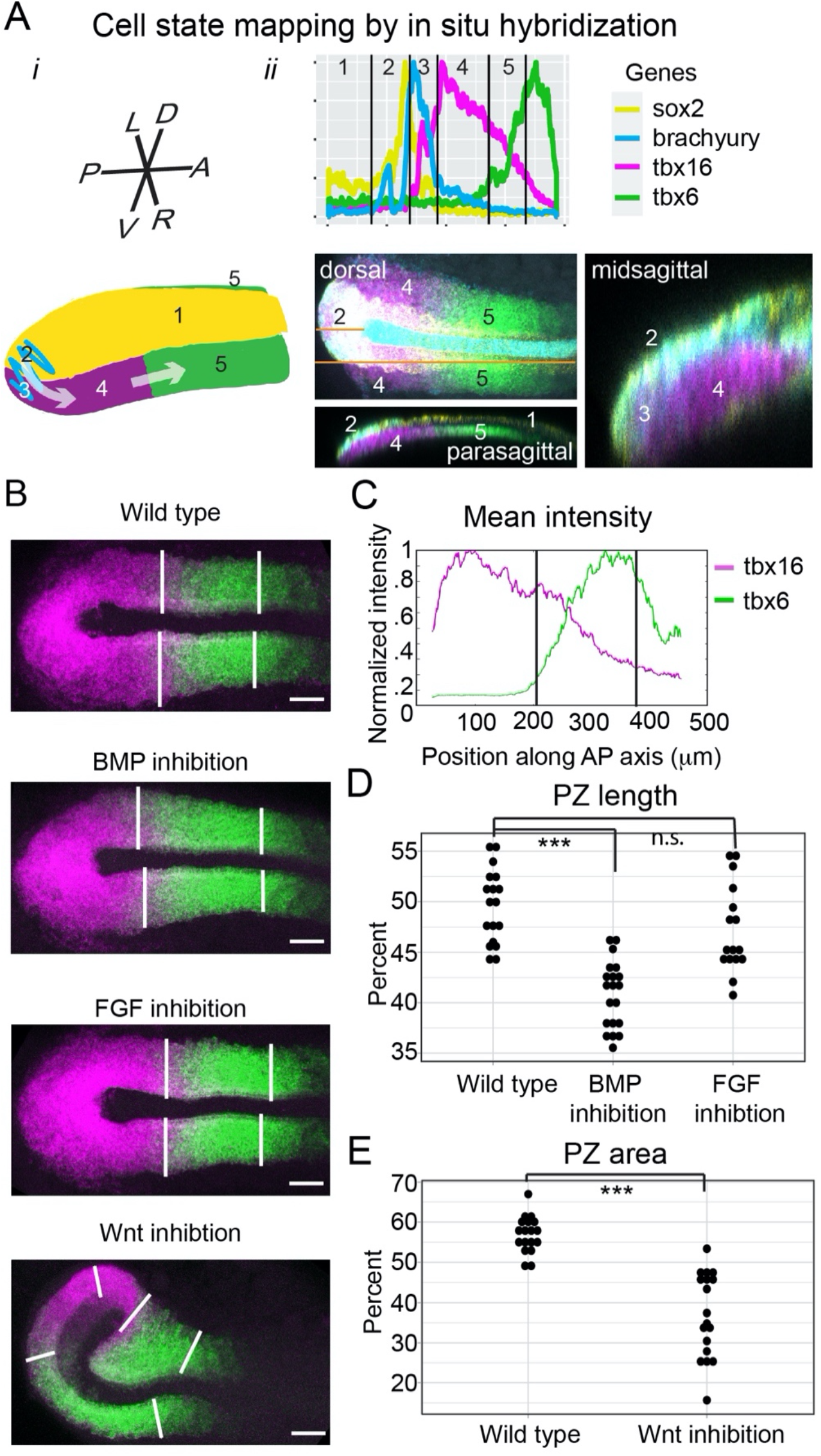
scRNAseq gene expression states map to the tailbud. **(A)** (*i*) A schematic showing the developmental trajectory of the paraxial mesoderm in the tailbud. All panels show the expression of *sox2* (yellow), *brachyury* (cyan), *tbx16* (magenta), and *tbx6* (green). (*ii*) Fluorescent in situ hybridization maps the transcriptional states (numbered) defined by scRNAseq onto the tailbud. In the dorsal view, the orange lines mark locations of the midsagittal (short line) and parasagittal (long line) slices. **(B)** Expression of *tbx16* (magenta) and *tbx6* (green). Vertical lines mark the transition from PZ to PSM and PSM to anterior PSM. Scale bar is 50 microns. **(C)** Plot of signal intensity in a representative wild-type embryo along the anterior-posterior axis. Vertical bars are cutoffs for the PZ and PSM of 20% and 85% of maximum *tbx6* expression, respectively. **(D)** PZ length normalized to total length of PZ and PSM in wild-type, BMP and FGF inhibited embryos. **(E)** PZ area normalized to total area in wild-type and Wnt inhibited embryos. *** is p<.001. See also Fig. S3.

To validate the scRNAseq analysis, we chose to test the predictions of changes in the abundance of neuronal and PZ states. First, the scRNAseq predicts that Wnt inhibited embryos would have more neuronal cells (Fig. 1E). This is consistent with reports that elimination of Wnt signaling leads NMPs to exclusively adopt a neuronal fate. In our milder perturbation of Wnt signaling, 1/3 of embryos have an abnormal cap of neuronal tissue covering the embryos’ posterior, confirming the scRNAseq results (Fig. S3).

A second prediction of the scRNAseq analysis is that the PZ is smaller in BMP and Wnt inhibited embryos but not in embryos subject to FGF inhibition. To test this prediction, we performed fluorescent in situ hybridization for a PZ marker, *tbx16,* and a PSM marker, *tbx6* (Fig 2B). In wild-type, BMP and FGF inhibited embryos, the *tbx16* and *tbx6* signal was measured along the anterior-posterior axis of the embryo for both the left and right sides (Fig. 2C). The PZ/PSM transition was set to the value derived from the scRNAseq analysis (20% of the maximum value of *tbx6*) and then the PZ length was normalized to the total tailbud length. Consistent with the scRNAseq analysis, BMP but not FGF inhibited embryos exhibited a decrease in PZ length (Fig. 2D). Due to the bent body axis exhibited by the majority of Wnt inhibited embryos, the area of the PZ and PSM were quantified. As predicted, Wnt inhibited embryos have a smaller PZ (Fig. 2E). Thus, this approach to analyzing scRNAseq data accurately identifies cell states that can be quantitatively mapped back onto the embryo.

### Cell motion states

We hypothesized that the same computational techniques used to classify gene expression states could be applied to cell motion data to objectively define the cell motion states (Fig. 3A). For this purpose, we used tracking data from confocal timelapse imaging of cells in the DM through PSM collected over 1-3 hours in wild-type embryos and embryos subject to signaling perturbations. As with the gene expression analysis, the cell motion statistics for each cell track were used to order the tracks in pseudotime, and the state transitions were defined using the change point detection algorithm. The cell states were color coded and spatially mapped back onto the embryo using the original cell track position. Initially, we chose not to use cell position as a pseudotime input both to facilitate pooling of data from multiple embryos together and to make the analysis analogous to that of the scRNAseq data which had all spatial information removed by cell dissociation. This procedure is successful solely using the statistics for cell velocity, average neighborhood cell speed within 20 micron radius of each cell, acceleration, and displacement over 6 and 15 minutes (Fig. 3B). The change point detection algorithm classifies the cells into four cell motion states (Fig. S4). These states are roughly segregated in space and their sequence matches the known developmental trajectory. Thus, cell migration states can be considered analogous to gene expression states.

**Fig. 3.**
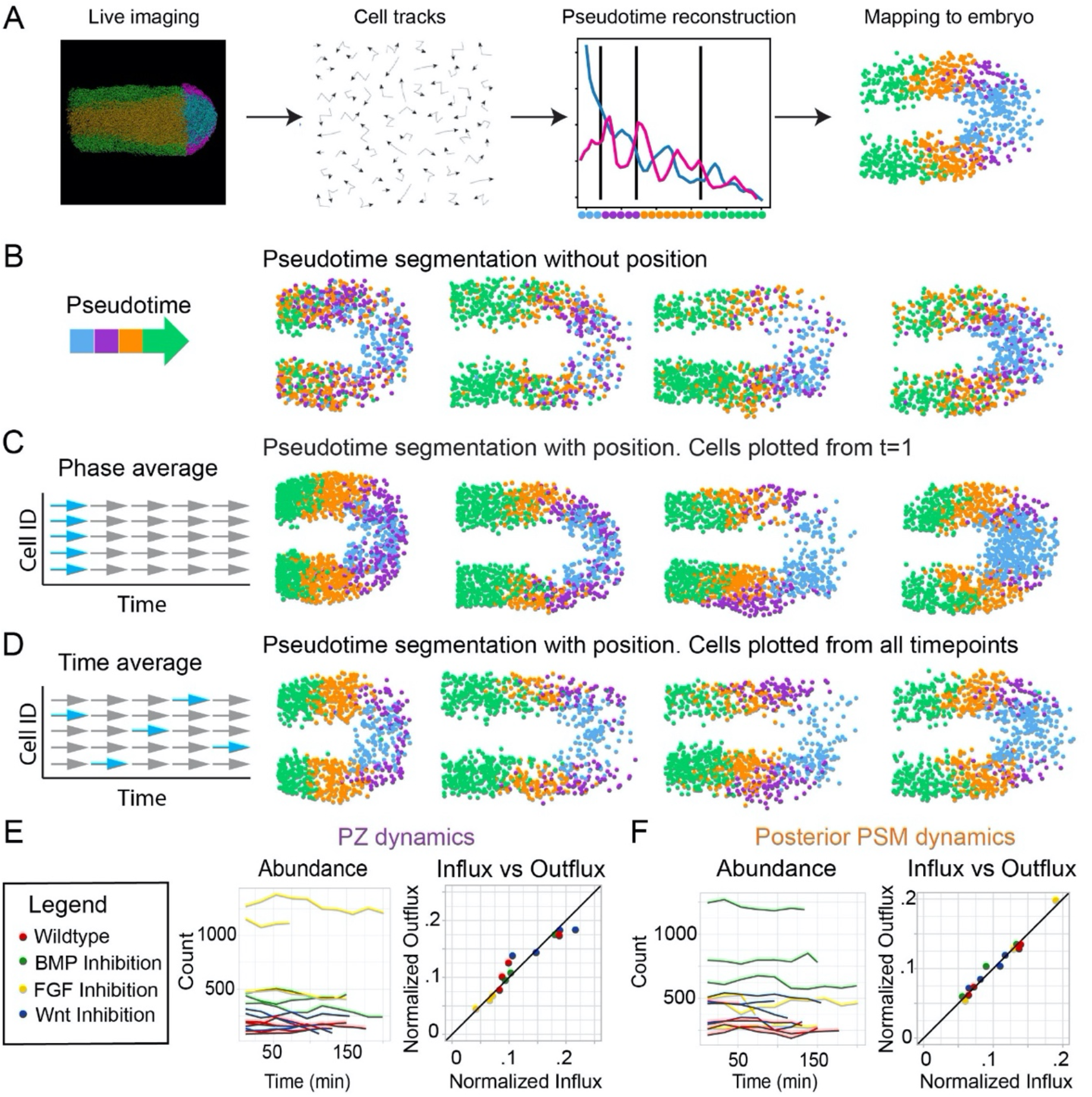
The pattern of cell motion states is ergodic. **(A)** Conceptual approach to identifying cell motion states. Tailbuds are imaged in 4D and cells are tracked, position is removed from cell tracks and data from multiple embryos are pooled, cell motion statistics used to construct and segment pseudotime, and cell states mapped back onto the embryo using the original cell position. **(B, C** and **D)** Cell motion state patterns of four wild-type embryos generated using three variations of this method. Each plot is a dorsal view with anterior orientated to the left. Each point is one cell, colored by migration state. (B) Plot of the segmentation of four embryos using a set of five cell motion parameters and data pooled from all embryos. (C and D) Cell state patterns generated individually for each embryo using cell position and cell track displacement for pseudotime estimation. For plotting, 1000 cells were chosen at random from each embryo from either the first timepoint (C) or from all timepoints (D). Note the similar patterns for each embryo, i.e. the vertically aligned cell state plots in C and D. **(E** and **F)** The PZ (E) and posterior PSM (F) cell counts over time and plots of cell influx vs outflux for each state. See also Fig. S4 and S5 and Movie S1.

### An ergodic pattern of cell motion states

The cell tracking data includes cell position, and we postulated that utilizing this information would improve the cell state segmentation. We therefore created a cell state map for each embryo using cell position and cell track displacement as inputs for pseudotime assembly (Fig. 3C, 3D and S5). These pseudotime sequences were then segmented based on the aforementioned cell motion statistics. This approach cleanly segmented the embryo into four cell states. As each embryo contains tens of thousands of data points, we plotted only a sample of the data from either a single time point, i.e. a phase average (Fig. 3C), or an identically sized selection randomly chosen from all time points, i.e. a time average (Fig. 3D). The distribution of states is extremely similar in both plots which is indicative of an ergodic system. The stability of the cell state pattern is evident in a movie generated using the average of each timepoint of our longest wild-type dataset (Movie S1). To obtain further evidence of ergodicity, we measured the cell abundance in each state over time as well as the influx and outflux from these states. The expectation is that the size of these domains would remain constant, and the fluxes would balance. We focused on the two states in the middle of the sequence, the PZ and posterior PSM, since we have both their complete influx and outflux data. Interestingly, the abundance and dynamics of these states can vary substantially from embryo to embryo and treatment to treatment, but the fluxes are balanced in each embryo (Fig. 3E and F). The balanced fluxes would help maintain the ergodic pattern of cell state transitions.

Given the stability of the migration state transitions, we wondered whether the transitions between gene expression states were also ergodic. Since RNA sequencing is an endpoint assay that does not readily lend itself to the calculation of time averages, we utilized RNA velocity, which considers the relative amount of intron and exon RNA for each gene, to estimate the flux between states. As expected, the overall RNA velocity is directed down the path of mesoderm differentiation (Fig. 4A and S6). Flux was calculated as the proportion of cells that transitioned to a different state (Fig. 4B and S7). For wild-type, the influx generally matches the outflux suggesting that the size of these domains is stable and that the patterns of cell state transitions may be ergodic. However unlike in the cell motion states, the balance between influx and outflux can be altered by perturbation of cell signaling.

**Fig. 4.**
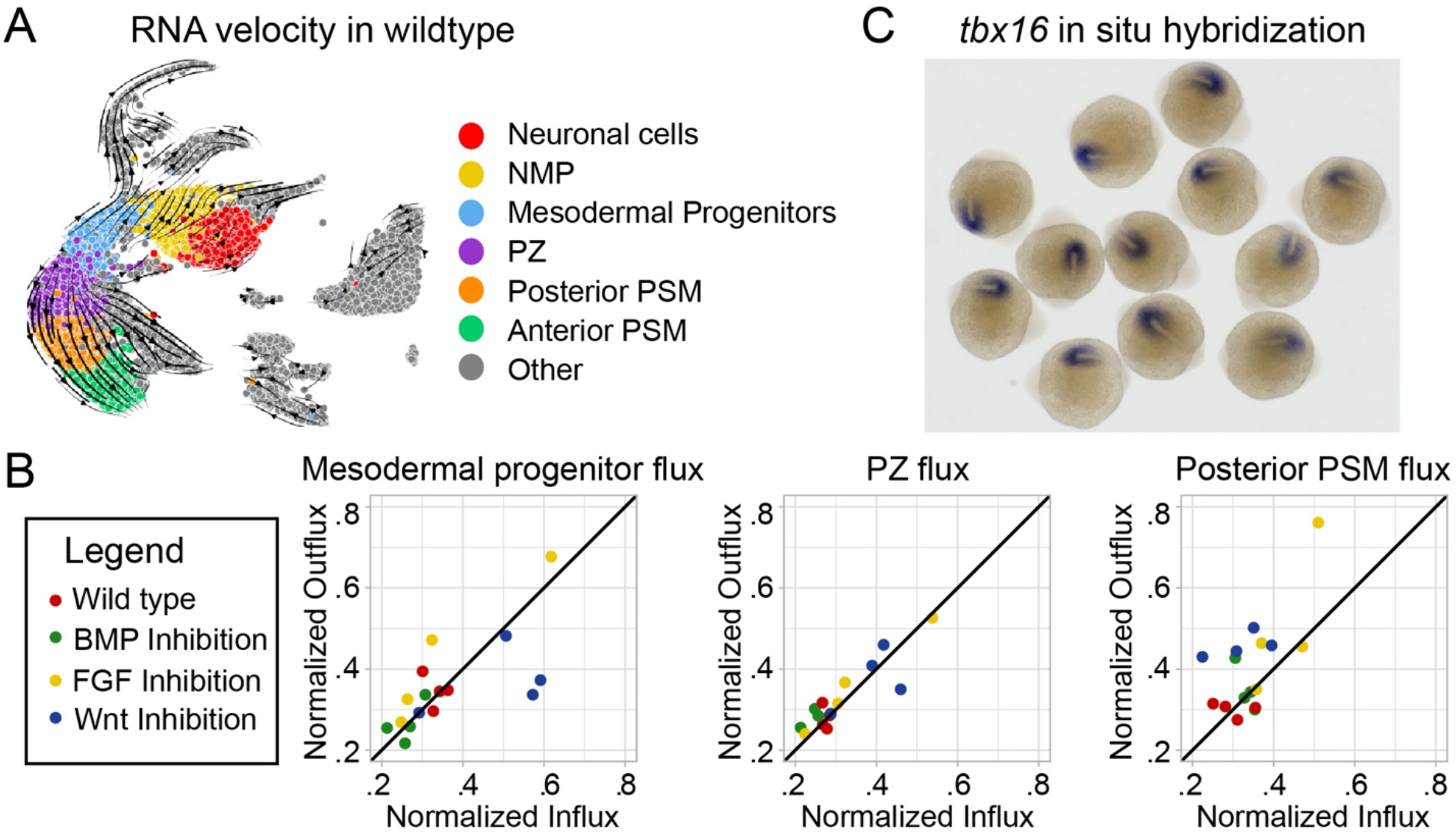
Flux analysis of gene expression cell states is consistent with ergodicity. **(A)** RNA velocity UMAP plot of tailbud gene expression states. **(B)** Influx vs outflux plots estimated by RNA velocity. Note that the influx and outflux are balanced in wild-type embryos, but cell signaling perturbation can alter this balance. **(C)** In situ hybridization for *tbx16* among sibling embryos illustrates the reproducibility of embryonic development. See also Fig. S6 and S7.

A batch of sibling embryos at roughly, but not exactly the same stage in development, produce very reproducible patterns of gene expression (Fig. 4C). In the tailbud, the consistency of this pattern is remarkable given the dynamics of cell motion that are driving elongation of the body axis. However, if the pattern of state transitions in cell motion is ergodic, it is not surprising that the gene expression patterns are also likely ergodic.

## Discussion

The ergodic pattern of cell state transitions may represent an emergent level of biological order that mediates gene network actuation of the stereotypical progression of embryogenesis. Our parallel analysis of gene expression and cell migration states using dimensional reduction and a change point detection algorithm demonstrates that these cell state transitions can be objectively defined and mapped back onto the embryo. While any time series dataset is well suited for an analysis of ergodicity, starting with cell state identification enables detection of ergodicity in complex datasets and reveals higher order ergodic patterns.

Ergodicity normally refers to a single stable state in which a dynamical system resides for a given amount of time. The length of time that a system remains in this state is referred to as the sojourn time. In this study, ergodicity refers to a pattern of successive cell states that remains stable over a period of 2-3 hours. Thus counterintuitively, this ergodicity does not mean that the tailbud doesn’t change, but that the pattern of changes doesn’t change. The ergodic pattern of cell state transitions could be thought of as a “meta-state”.

This biological ergodicity is a dynamic order that arises from the genome and the biochemical and physical interactions among cells in space and time. This ergodicity is dependent upon the length of the time interval being studied. If one were to combine data from a gastrula with data from an embryo during body elongation, then there would likely be no ergodicity. Thus, there is a sojourn time for a given pattern of cell state transitions that will scale with the developmental process under study. A given ergodic pattern may exhibit a sojourn time of hours in the case of the zebrafish tailbud or years in the case of adult homeostatic tissues. During development, the embryo may transition from one ergodic pattern to another as it develops until it reaches the relatively long sojourn time of homeostatic tissues in the mature organism.

An innovation of this study is the finding that cell position and cell motion statistics are sufficient to identify cell states via dimensional reduction and subsequent segmentation into cell states using the change point detection algorithm. These dimension reduction techniques, developed to analyze scRNAseq data, can be applied to other complex datasets along with the change point detection algorithm to identify underlying patterns. The methodology presented here provides a way to assay the validity of the assumption of ergodicity, to identify experimental conditions in which ergodicity is lost, and to measure the time intervals over which ergodicity is maintained in complex datasets. For example, homeostasis should be congruent with ergodicity, and a breakdown of homeostasis due to mutation, aging or disease could be quantified via an analysis of ergodicity.

*C. elegans* embryos are famous for their invariant cell lineages in that one embryo develops in exactly the same manner as any other *C. elegans* embryo. Vertebrate embryos do not display these invariant cell lineages, but fate mapping demonstrates that subpopulations of cells reproducibly give rise to specific tissues in every embryo of a given species. Thus, while *C. elegans* development is precisely reproducible down to the cellular level, vertebrate embryonic development is reproducible down to the level of ensembles of cells. The question is how is this reproducibility achieved in vertebrate embryos? Some of the reproducibility of development is due to gene networks that specify and maintain quasi-stable states through which cells transit during development. For example, neuromesodermal progenitors transition to mesodermal progenitors, then to presomitic mesoderm and then to somites. It follows that in vertebrate embryos, these gene regulatory networks operate at the level of ensembles of cells as reflected in the concepts of developmental regulation and community effect. This study finds that the pattern of these transitory cell states is ergodic and therefore dynamically stable over time. The fact that the pattern of cell state transitions doesn’t change indicates that the rates of change are stable. The absolute cell state influxes and outfluxes vary significantly between embryos but are balanced in each embryo. Thus, ergodicity exists at a higher level, the derivative, and may represent an emergent systems-level order linking gene regulatory networks with the general reproducibility of embryonic development.

## Acknowledgements

We thank Abdel-Rahman Hassan for thoughtful discussions, Sarah Smith, A-R Hassan, Abby Kindberg and Holger Knaut for comments on the manuscript, Guilin Wang, Mei Zhong, the Yale Keck Biotechnology Resource Laboratory and the Yale Stem Cell Genomics Core Facility for help with the scRNAseq.

## Funding

National Institute of Health grant 1F32GM137502-01 (MAG)

National Institute of Health grant R01GM129149 (SAH)

### Author Contributions

MAG performed wet lab experiments, data analysis, data interpretation and wrote the manuscript, YK led the data analysis, performed data interpretation and wrote the manuscript, DJ contributed to the wet lab experiments, HK interpreted data and supervised data analysis, and SAH conceived of and supervised the project, interpreted data and wrote the manuscript.

## Materials and Methods

### Data and code availability

The scRNAseq data has been archived at NCBI GEO (accession no: GSE173894).

### Zebrafish methods

Tüpfel-longfin zebrafish were raised according to standard protocols approved by the Institutional Animal Care and Use Committee. Experiments were performed before sex determination in zebrafish. FGF, BMP, and Wnt signaling perturbations were performed using protocols previously developed to modulate cell migration. Specifically, starting at the 6-somite stage embryos were incubated in 50 mM of SU5402 or 40 mM of DMH1 for two hours to inhibit FGF or BMP signaling, respectively. Wnt signaling was inhibited by injecting *notum-1* mRNA at a concentration of 150 ng/mL into embryos at the single cell stage and then incubating them until the 10-somite stage. This treatment yields a phenotypic spectrum, and embryos with nascent body elongation defects were chosen for further experiments.

### Tailbud Dissections and scRNA sequencing

Embryos were incubated until the 10-12 somite stage and then dissected in ice cold Hank’s Balanced Salt Solution. The tail was collected by cutting immediately posterior to the last formed somite. Groups of tails consisting of ten tails for wild-type, FGF, or BMP inhibition or twelve tails for Wnt inhibition were pooled together. Cells were dissociated by incubation in 20 U/mL papain solution (Worthing Biochemical) for 15 minutes at 29 °C with gentle agitation. Halfway through the incubation the solution was triturated ten times with a P200 pipette. Cells were spun down at 300g for five minutes and then resuspended in 40 mL of cold HBSS. Cell concentration and viability were checked with a hemocytometer and the volume of the solution was adjusted if required.

#### Construction of 10X Genomic Single Cell 3’ RNA-Seq libraries (Version 3) and sequencing with an Illumina HiSeq4000

##### GEM Generation and Barcoding

Single cell suspension in RT Master Mix was loaded on the Single Cell A Chip and partition with a pool of about 750,000 barcoded gel beads to form nanoliter-scale Gel Beads-In-Emulsions (GEMs). Each gel bead has primers containing (i) an Illumina R1 sequence (read 1 sequencing primer), (ii) a 16 nt 10x Barcode, (iii) a 10 nt Unique Molecular Identifier (UMI), and (iv) a poly-dT primer sequence. Upon dissolution of the Gel Beads in a GEM, the primers are released and mixed with cell lysate and Master Mix. Incubation of the GEMs then produces barcoded, full-length cDNA from poly-adenylated mRNA.

##### Post GEM-RT Cleanup, cDNA Amplification and library construction

Silane magnetic beads were used to remove leftover biochemical reagents and primers from the post GEM reaction mixture. Full-length, barcoded cDNA was then amplified by PCR to generate sufficient mass for library construction. Enzymatic fragmentation and size selection were used to optimize the cDNA amplicon size prior to library construction. R1 (read 1 primer sequence) were added to the molecules during GEM incubation. P5, P7, a sample index, and R2 (read 2 primer sequence) were added during library construction via End Repair, A-tailing, Adaptor Ligation, and PCR. The final libraries contain the P5 and P7 primers used in Illumina bridge amplification.

##### Sequencing libraries

The Single Cell 3’ Protocol produces Illumina-ready sequencing libraries. A Single Cell 3’ Library comprises standard Illumina paired-end constructs which begin and end with P5 and P7. The Single Cell 3’ 16 bp 10x Barcode and 10 bp UMI are encoded in Read 1, while Read 2 is used to sequence the cDNA fragment. Sequencing a Single Cell 3’ Library produces a standard Illumina BCL data output folder. The BCL data includes the paired-end Read 1 (containing the 16 bp 10x Barcode and 10 bp UMI) and Read 2 and the sample index in the i7 index read.

### Preprocessing of scRNA sequencing data

We aligned the scRNA-seq data to Grcz11 and demultiplexed using Cell Ranger (10X Genomics). After the generation of expression matrices for each sample, we utilized Seurat v3 for preprocessing and clustering of scRNA-seq data. First, we excluded cells with an ectopic number of genes or exceeding a specified percentage of mitochondrial genes (Table S1) based on visual inspection for the distribution of these statistics. After the filtering genes, we conducted integration following Seurat's SCTransform integration.

We applied principal components analysis and embedded the 30-dimensional PCA coordinates into 2 dimensional UAMP. We clustered cells by Seurat function “FindClusters” with a resolution parameter of 0.5.

### Pseudotime estimation of scRNA-seq

To recover cell state dynamics encoded in the gene expression data, we ordered a subset of scRNA-seq cells which belong to the axis from ADM to PSM so that its ordering recapitulates the developmental trajectory during body elongation. In particular, we embedded the z-scores of 30 dimensional PCA coordinates of cells belonging to specified clusters (Sox3+, Sox2+, DM, PZ, pPSM and aPSM) into one dimensional UMAP coordinates. Here, we expected that the most variable axis within gene expression space during this process would be the developmental trajectory. For UMAP embedding, we used the “umap-learn” package in Python and set “n_neighbors” as 400 and “min_dist” as 0.1.

### Segmentation of scRNA-seq pseudotime

We segmented the pseudotime trajectory of scRNA-seq into several segments within which each cell *c* has similar z-scores of 30-dimentional PCA coordinate *x*_*c*_ in order to dissect the dynamics along the progression of cell state transitions during zebrafish body elongation. We utilized a Bayesian algorithm of change detection to find break points of segments *b*_*k*_(*k* = 1, ..., *K*) which minimize the total error from the mean of profile of the segment ∑_*k*_ *E*_*k*_ where 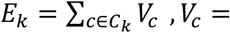 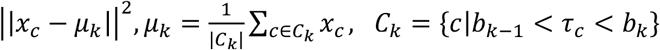 and *τ*_*c*_ is the discretized rank of the estimated pseudo time. We discretized the pseudotime rank into 30 bins for computational efficiency. We determined K as 5 scRNA-seq data using the elbow method which chose a saturation point along the group variation curve as function of the number of clusters.

### Flux analysis based on RNA velocity

We recovered cell state dynamics behind scRNA-seq data using scVelo which estimates the velocity of RNA for each single cell Using a computed velocity *v*_*c*_ of cell *c* with its single cell transcriptome *x*_*c*_, we calculated the predicted transcriptome after a micro duration *δ* as *x*′_*c*_ = *x*_*c*_ + *δv*_*c*_. We set *δ* so that 3% of transcriptome *x*_*c*_ changed during *δ*. We conducted PCA analysis on a concatenated expression matrix of current and *δ*-elapsed transcriptome and used the 10-dimensional PCA coordinates of cell *c* at current and *δ*-elapsed time points, which we denoted as *z*_*c*_ and *z*′_*c*_ We estimated the segment *b*′_*c*_ which cell *c* after *δ* is belonging to as the current segment *b*_*c′*_ of the cell *c*′ whose PCA coordinates *z*_*c′*_ are the nearest to the predicted *δ*-elapsed PCA coordinates *z*′_*c*_. We quantified the transition 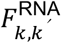 from segment *k* to segment *k*′ as the count of cells whose current and *δ*-elapsed segments are *k* and *k*′ respectively. The normalization is done for the total number of cells within the segment. We also defined the influx rate and outflux rate of segment *k* as the normalized summation of transition from any segments to *k* and from *k* to any segments and defined them as 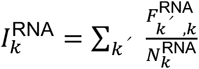 and 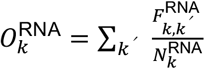 respectively, where 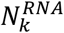 is number of cells in segment *k*.

### Estimation and segmentation of cell movement pseudotime

We recovered the pseudo dynamics of the cell movement properties by pseudotime estimation followed by variance minimization segmentation. For pseudotime estimation without positional information, we pooled together the embryos and calculated 1-dimentional UMAP embeddings of each cell from the z-scores of its speed, acceleration, magnitude of neighborhood velocity, and displacement distance for 6 minutes and 15 minutes. The magnitude of neighbor velocity is defined as ||∑_*c∈N(c)*_ *v*_*c*_|| where *N*(*c*) = {*c*′| ||*p*_*c*_ − *p*_*c′*_|| < 20μ*m*} where *p*_*c*_ and *v*_*c*_ are position and velocity of cell *c* and ||.|| is the Euclidean norm. For the pseudotime estimation with positional information each embryo was processed separately. We embedded the z-scores of 3D spatial position and 3D displacement during 15 minutes into one dimensional order using UMAP. For UMAP embedding, we used the “umap-learn” package in Python and set “n_neighbors” as 100 and “min_dist” as 0.1. We segmented both types of cell movement pseudotimes using the same methodology for the segmentation of scRNA-seq pseudotime, except that the properties *x*_*c*_ for minimizing within-group variation are the z-scores of the speed, acceleration, magnitude of neighborhood velocity, and displacement distance for 6 minutes and 15 minutes. We specified the number of change points *K* as 3 using the elbow method which chose a saturation point of along the group variation curve as function of the number of clusters.

### Flux analysis between cell movement segments

We quantified the flux between segments in cell movement data utilizing cell tracking information. We counted the cells which stay at segment *k* at a time point *t* and transit to segment *k*′ at the subsequent time point *t* + 1. In the same way as the previous section, we defined influx rate and outflux rate of segment *k* as the normalized summation of transition from any segments to *k* and that from *k* to any segments and defined them as 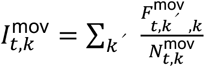 and 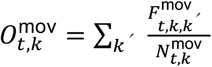 respectively, where 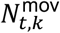is the number of cells in segment *k* at time *t*.

### Multicolor fluorescent in situ hybridization

Probes for *sox2*, *brachyury, tbx16,* and *tbx6* were purchased from Molecular Instruments. The hairpins and colors are listed in the table below. Staining of 10-12 somite embryos was performed using their recommended protocol with a few modifications. Specifically, batches of 15 embryos were stained simultaneously. The *tbx6* probe was diluted 1:10 to avoid excessive bleed through into the *sox2* channel. DAPI was added to the amplification mixture. After staining embryos were taken through a series of 25%/50%/75% glycerol in PBS. The posterior half of the embryo was isolated and mounted dorsal side up in 75% glycerol. Embryos were imaged with a Zeiss LSM 880 Airyscan Confocal using a 20x objective.

| Gene | Hairpin | Dye |
| --- | --- | --- |
| Sox2 | B1 | Alexa Fluor 546 |
| Tbx16 | B2 | Alexa Fluor 647 |
| Brachyury | B3 | Alexa Fluor 488 |
| Tbx6 | B4 | Alexa Fluor 594 |

Preprocessing of the microscopy images was done using ImageJ. The *sox2* and *tbx6* channels were subtracted from each other to eliminate bleed through. Images were rotated to a consistent orientation and a max intensity projection was created. Adaxial cells were identified in the DAPI channel and manually removed from the image. The midline separating the embryo into left and right halves was identified manually. Subsequent quantification was performed in Matlab. The image was smoothed with a Gaussian filter, and the region of interest was thresholded using Otsu’s algorithm on both the *tbx16* and *tbx6* channels. For wildtype, BMP, and FGF inhibited embryos, average fluorescent intensity was measured along the x axis of the image and normalized to the maximum value. This was done separately for the left and right sides of the embryo. The PZ/PSM boundary was taken to be the first point with a value greater than 20% of the maximum *tbx6* value. The anterior end of the PSM was defined as the last point greater than 85% of the maximum *tbx6* value. The scaled PZ length was the PZ length divided by the distance from the end of the tail to the anterior boundary of the PSM.

Wnt inhibited embryos had some modifications to the quantification. In bent embryos, the boundary separating the left and right halves was taken to be a line through the midpoint of the tailbud *brachyury* signal to the notochord and then following the notochord towards the head. For wild-type and Wnt inhibited embryos the outer perimeter of the embryo was traced manually. The curve was smoothed with a Savitzky-Golay filter and defined as the embryo’s axis. Pixels in the ROI were mapped to their closest points on the perimeter using the distance2curve function from John D’Errico. The mean intensity along the axis was calculated using a sliding window. The same thresholds were used for the PZ and PSM as described previously. The boundaries for these regions were taken to be a perpendicular dropped from the axis at the cutoff point. The scaled PZ area was the PZ area divided by the area of the PZ plus PSM.

Statistics were calculated using Mann-Whitney’s U.

---

**Figure S1.**
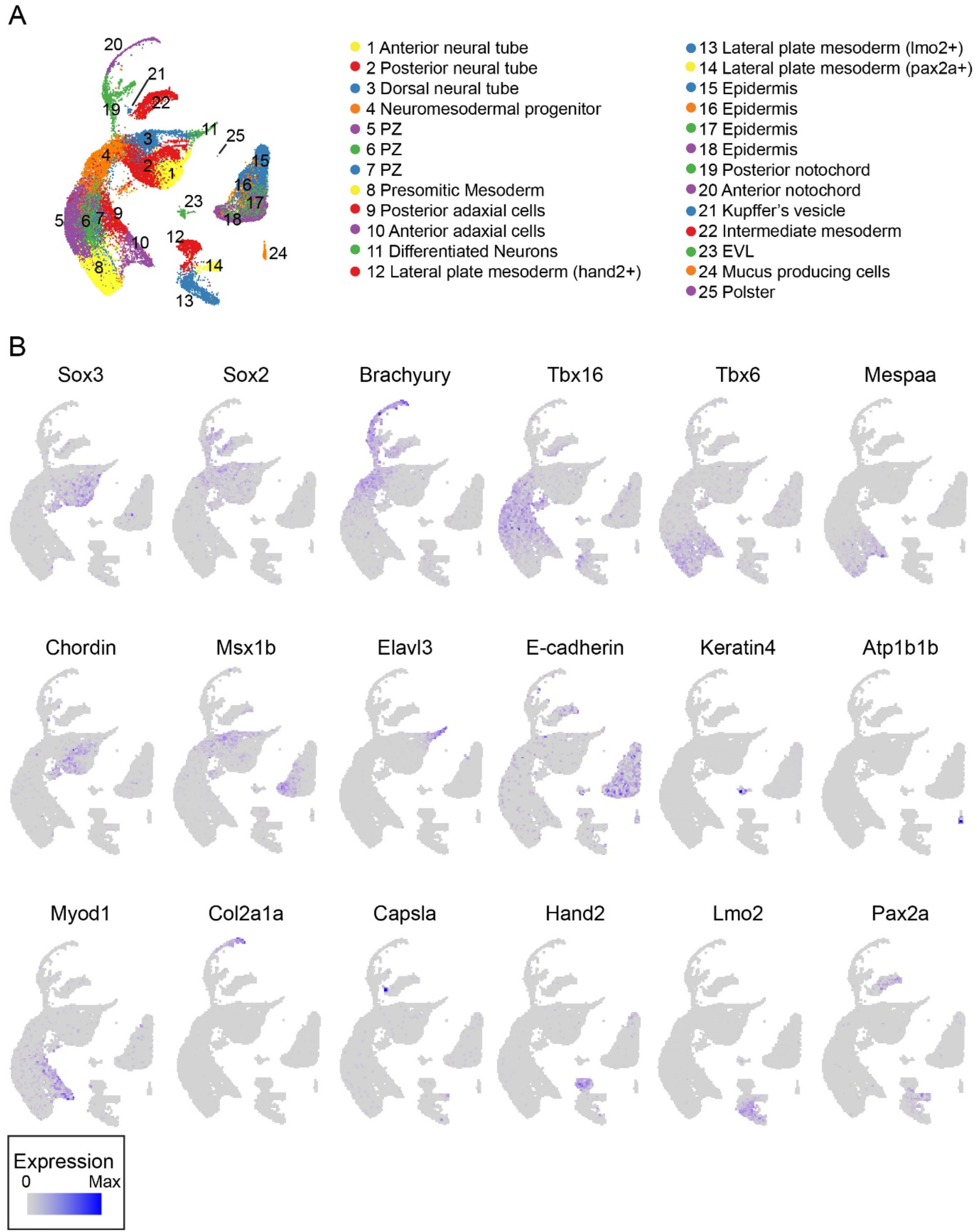
Gene expression states from scRNAseq. Related to Figure 1. **(A)** UMAP projection of cell clusters defined by Louvain clustering using Seurat. Clusters were manually annotated using marker genes. **(B)** UMAP projection of selected marker genes used to identify cell types.

**Figure S2.**
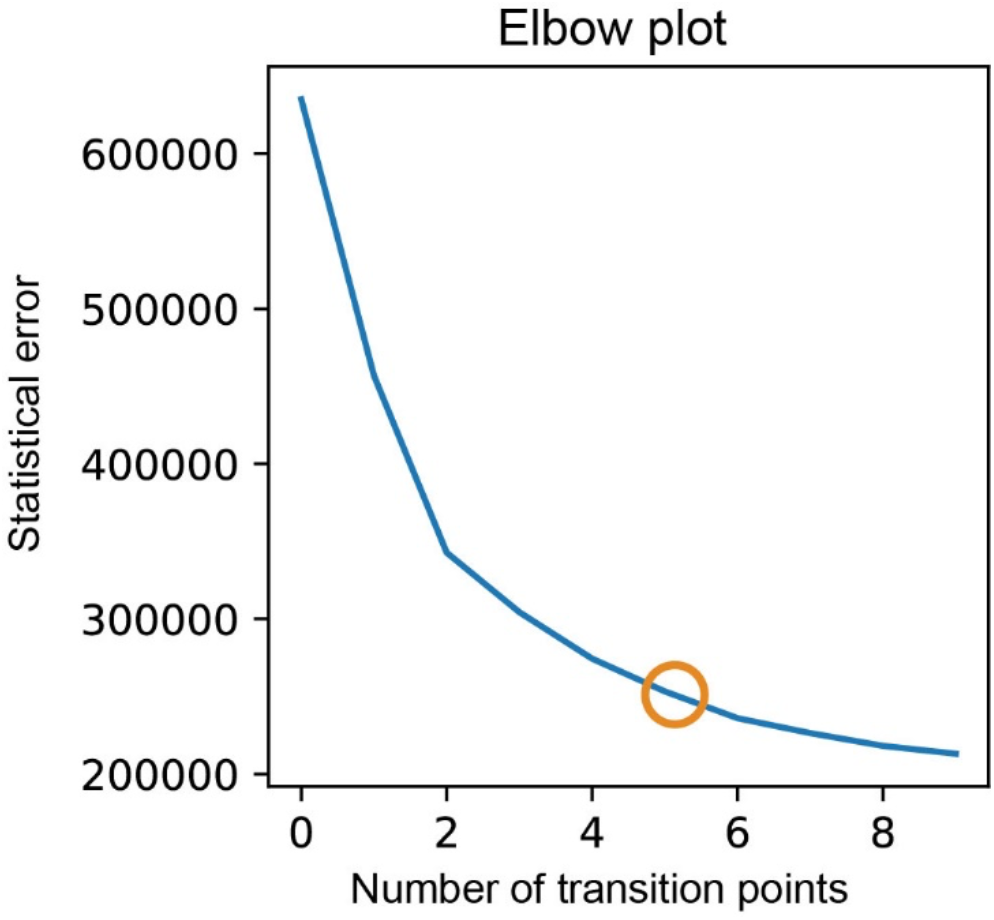
Segmentation of gene expression pseudotime. Related to Figure 1. An elbow plot used to select the number of transition points in the scRNAseq pseudotime. Circle marks the chosen number of transitions.

**Figure S3.**
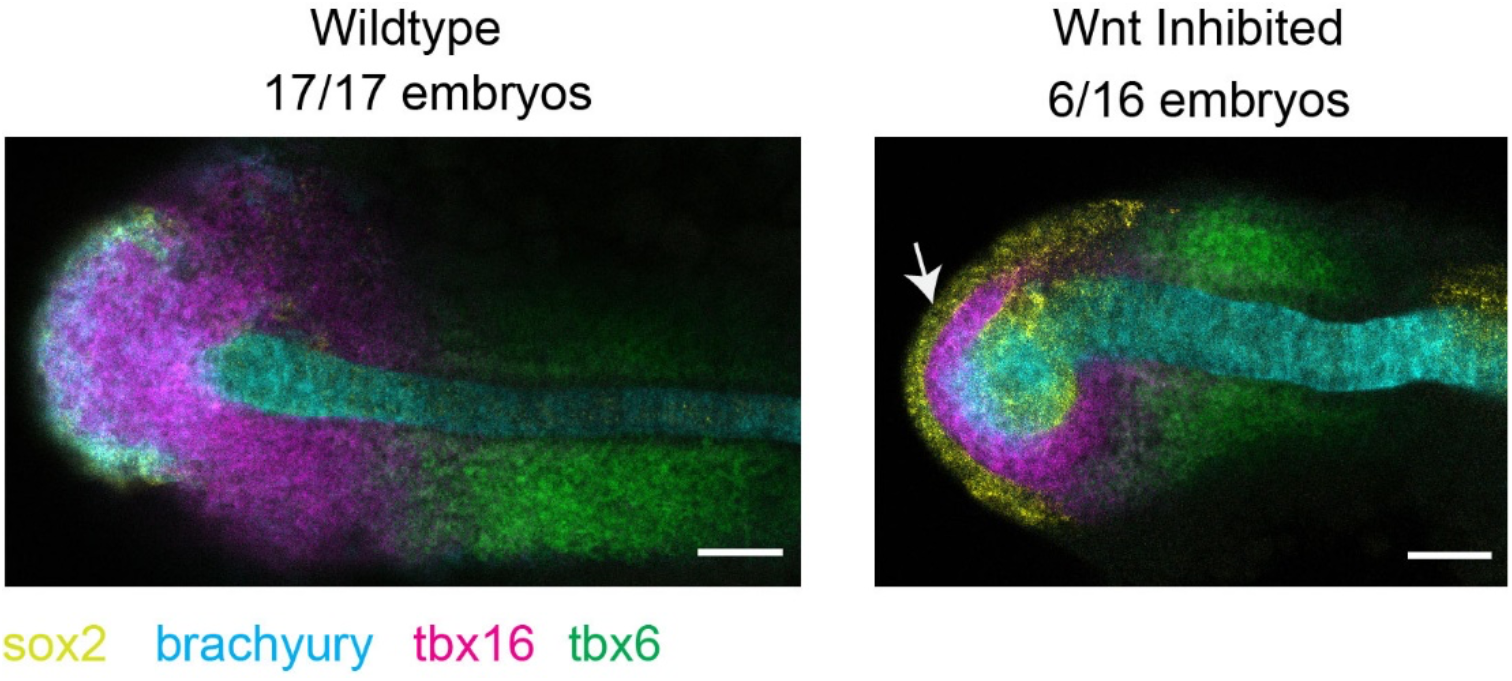
Wnt inhibited embryos have excessive neural tissue. Related to Figure 2. Five-micron thick projections through the ventral tailbud of a multicolor fluorescent in situ hybridization. Arrow points to inappropriately located *sox2* single positive neural tissue. Scale bar= 50 microns

**Figure S4.**
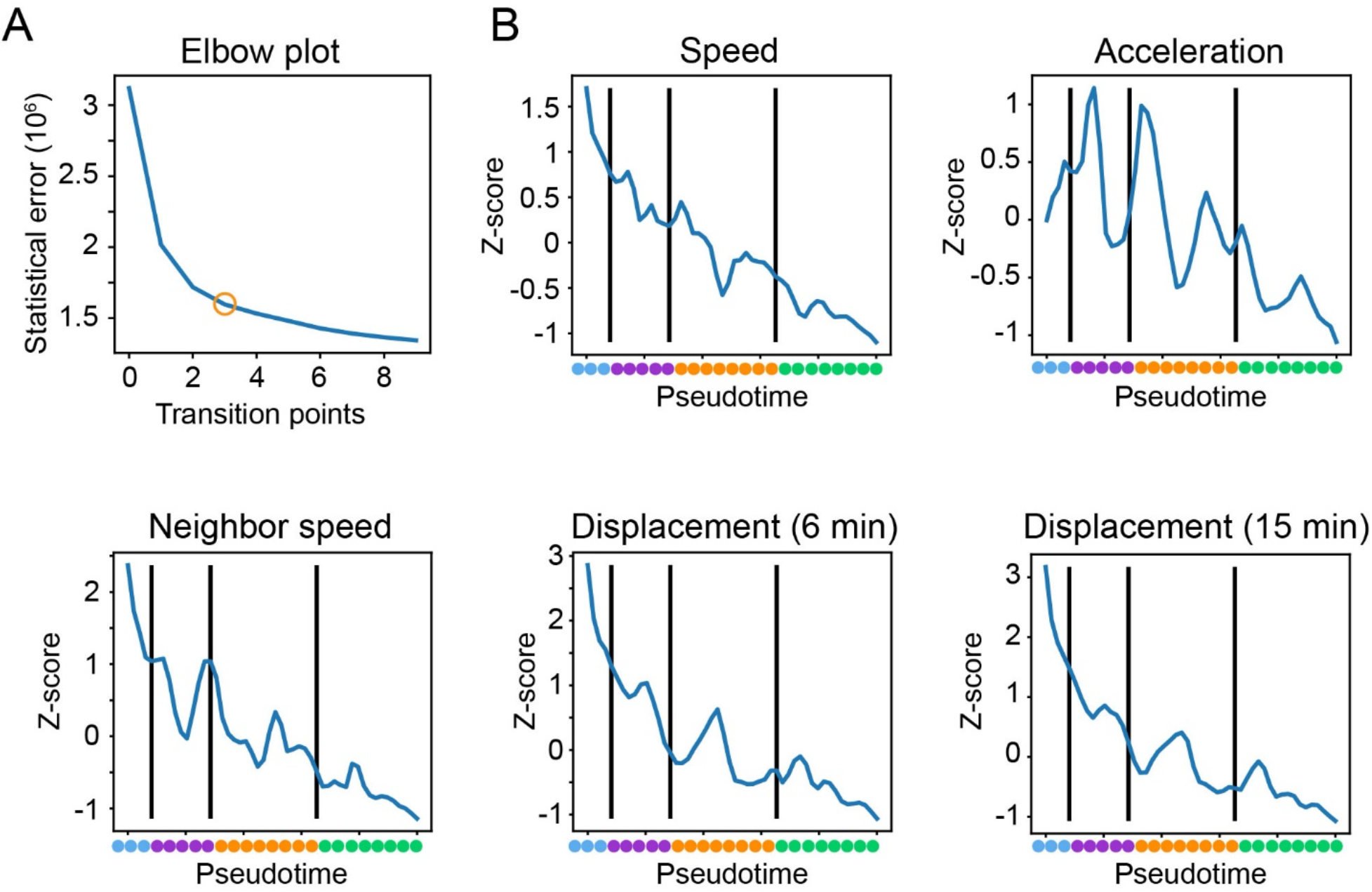
Cell motion pseudotime segmentation. Related to Figure 3. **(A)** An elbow plot used to select the number of transition points in the cell motion pseudotime. Circle marks the chosen number of transitions. **(B)** Cell motion statistics used to segment the embryos in Fig 3B plotted over pseudotime. Neighbor speed represents the average cell speed withing a 20 μm radius of each cell. Displacement statistics were analyzed over two time intervals. Vertical lines mark transition points. The segment colors along the pseudotime axis correspond to the cell motion state colors in Fig. 2 and S5.

**Figure S5.**
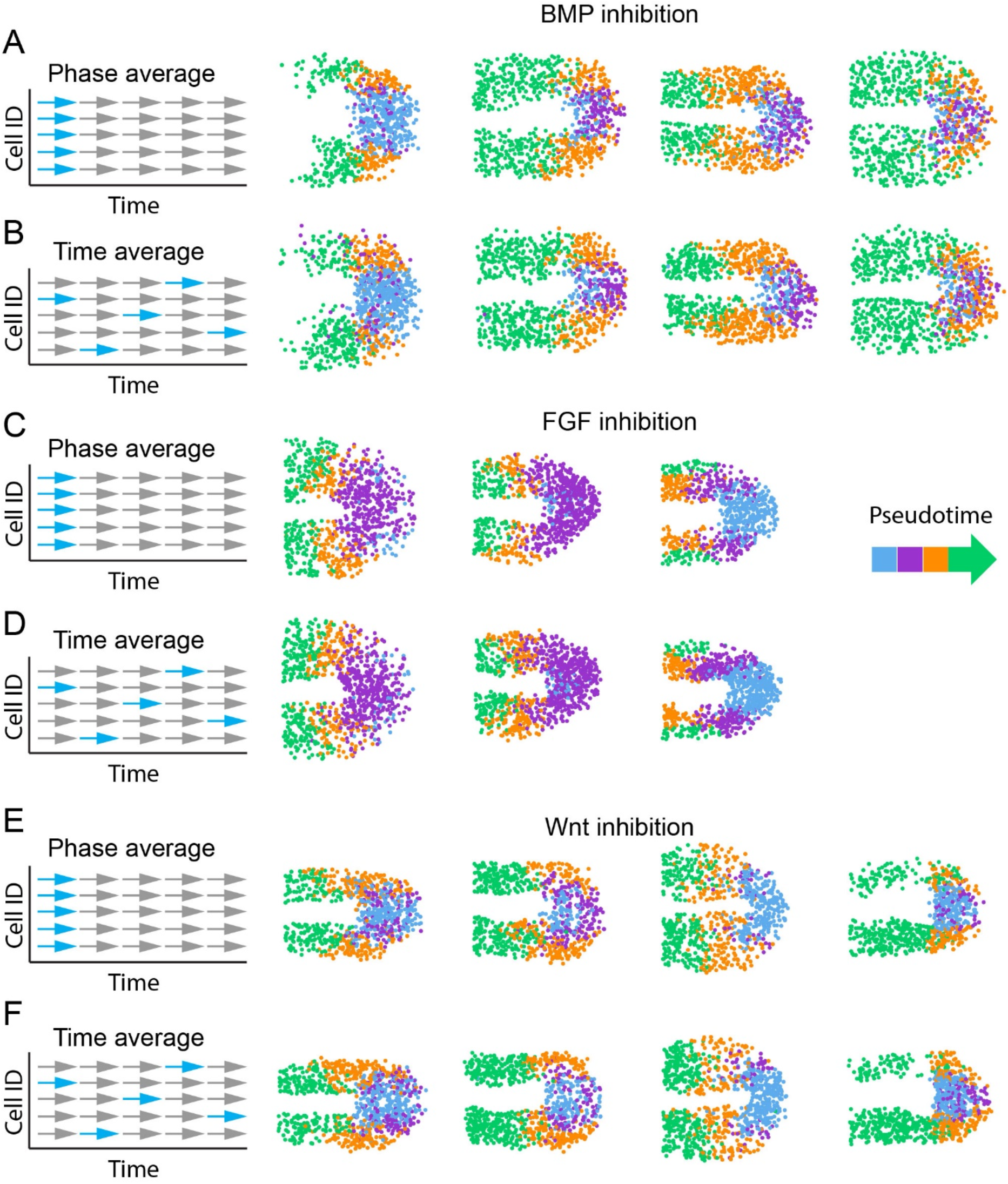
The pattern of cell migration states remains ergodic in embryos subject to signaling perturbations. Related to Figure 3. Data from embryos subject to signaling perturbations, processed and plotted identically to the wildtype embryos in Fig 3C. **(A)** The phase average plots for four embryos with reduced Bmp signaling. **(B)** Time average plots for the same four embryos with reduced Bmp signaling. **(C)** The phase average plots for three embryos with reduced FGF signaling. **(D)** Time average plots for the same three embryos with reduced FGF signaling. **(E)**The phase average plots for four embryos with reduced Wnt signaling. **(F)** Time average plots for the same four embryos with reduced Wnt signaling.

**Figure S6.**
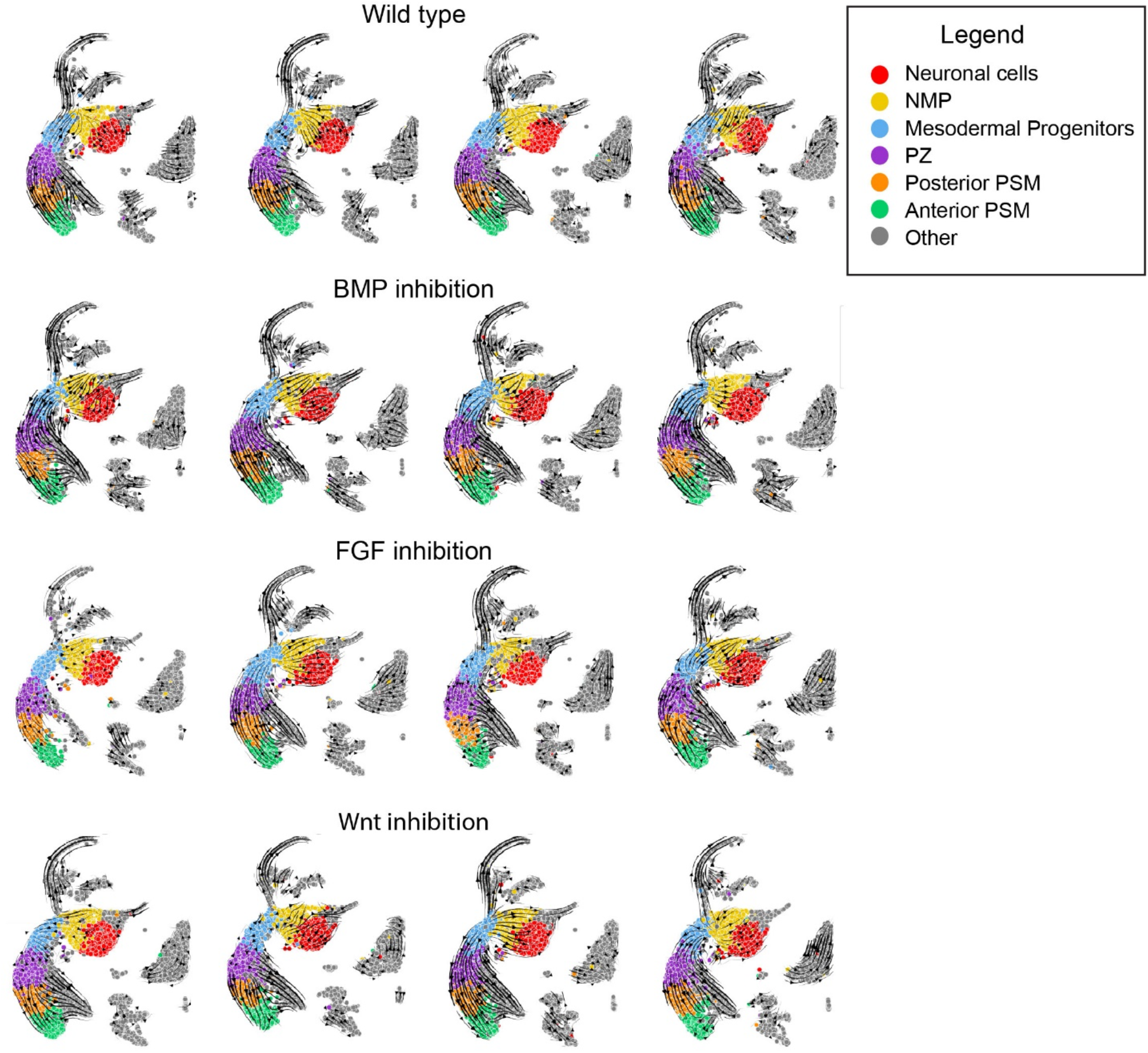
RNA velocity plots of all scRNAseq replicates. Related to Figure 4. Plots for all replicates were used to calculate cell state flux in Fig 4B.

**Figure S7.**
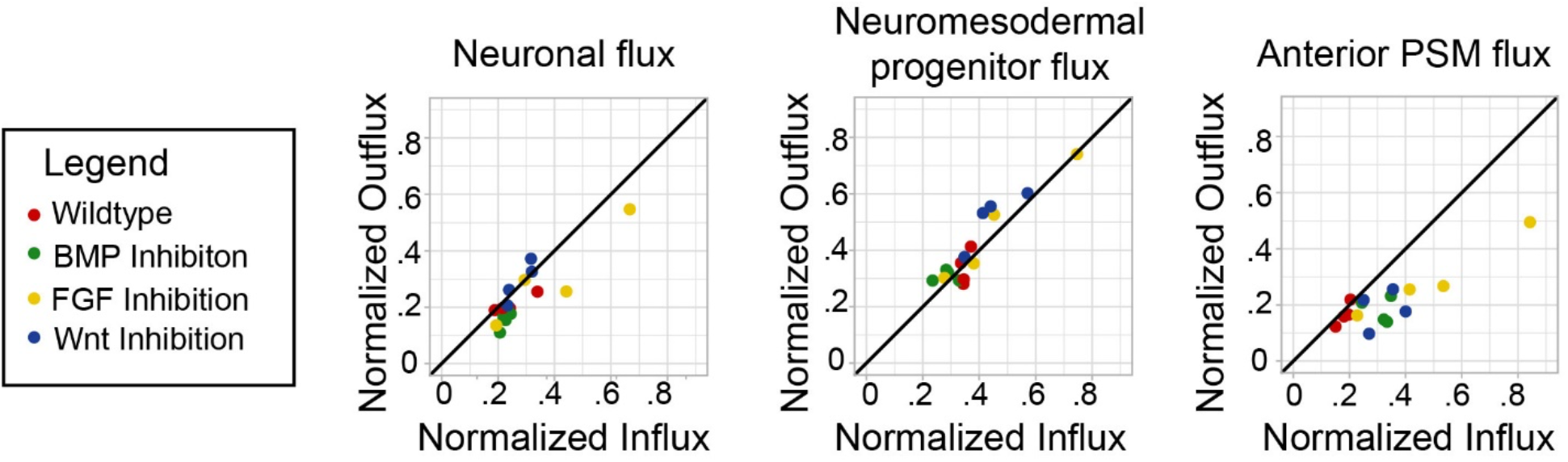
Cell flux through tailbud cell states derived from RNA velocity. Related to Figure 4. Influx is greater than outflux for the anterior PSM and neuronal states. However, this may be an artifact of the absence of further differentiated tissues in the dataset.

**Movie S1.**
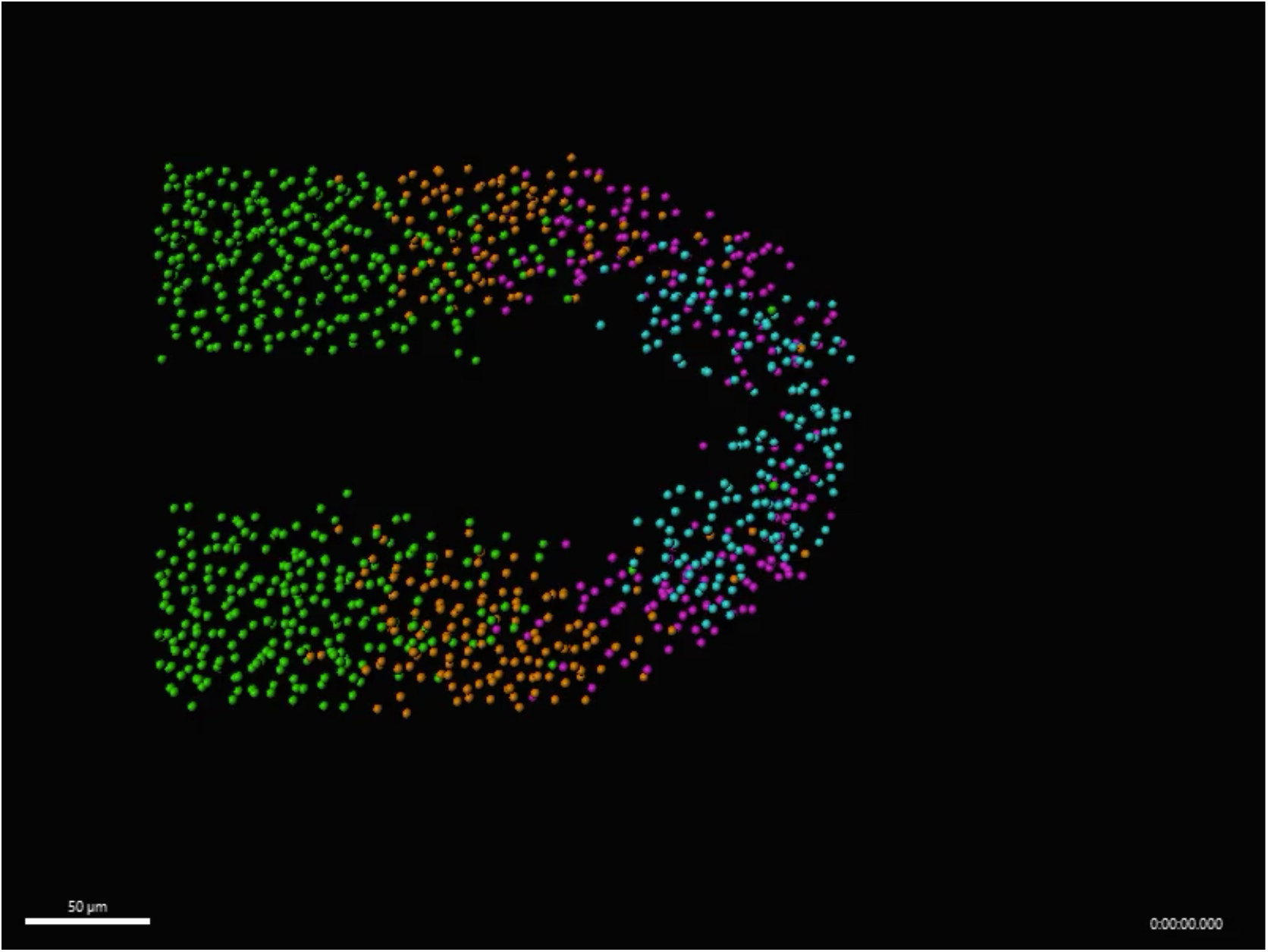
Ergodic pattern of cell state transitions. Related to Figure 3. A timelapse was generated using the phase average for each timepoint in the longest wild-type dataset. The states were segmented using position and displacement.

**TableS1.** Filtering criterions for the cells of scRNA-seq in each sample. Related to Figure 1.

| Samples | Max gene number | Minimum gene number | Max mitochondrial percentage |
| --- | --- | --- | --- |
| WT1 | 2800 | 1200 | 5 |
| WT2 | 2000 | 800 | 5 |
| WT3 | 2500 | 800 | 5 |
| WT4 | 2500 | 800 | 5 |
| BMP1 | 4000 | 1000 | 7 |
| BMP2 | 3000 | 1000 | 7 |
| BMP3 | 4000 | 1000 | 7 |
| BMP4 | 4000 | 1000 | 7 |
| FGF1 | 2000 | 800 | 10 |
| FGF2 | 2500 | 800 | 7 |
| FGF3 | 3000 | 800 | 7 |
| FGF4 | 4000 | 1500 | 7 |
| WNT1 | 2500 | 500 | 3 |
| WNT2 | 2500 | 500 | 3 |
| WNT3 | 3000 | 500 | 3 |
| WNT4 | 3750 | 500 | 3 |

